# Intranasal parainfluenza virus-vectored vaccine expressing SARS-CoV-2 spike protein of Delta or Omicron B.1.1.529 induces mucosal and systemic immunity and protects hamsters against homologous and heterologous challenge

**DOI:** 10.1101/2024.09.12.612598

**Authors:** Hong-Su Park, Yumiko Matsuoka, Celia Santos, Cindy Luongo, Xueqiao Liu, Lijuan Yang, Jaclyn A. Kaiser, Eleanor F. Duncan, Reed F. Johnson, I-Ting Teng, Peter D. Kwong, Ursula J. Buchholz, Cyril Le Nouën

## Abstract

The continuous emergence of new SARS-CoV-2 variants requires that COVID vaccines be updated to match circulating strains. We generated B/HPIV3-vectored vaccines expressing 6P-stabilized S protein of the ancestral, B.1.617.2/Delta, or B.1.1.529/Omicron variants as pediatric vaccines for intranasal immunization against HPIV3 and SARS-CoV-2 and characterized these in hamsters. Following intranasal immunization, these B/HPIV3 vectors replicated in the upper and lower respiratory tract and induced mucosal and serum anti-S IgA and IgG. B/HPIV3 expressing ancestral or B.1.617.2/Delta-derived S-6P induced serum antibodies that effectively neutralized SARS-CoV-2 of the ancestral and B.1.617.2/Delta lineages, while the cross-neutralizing potency of B.1.1.529/Omicron S-induced antibodies was lower. Despite the lower cross-neutralizing titers induced by B/HPIV3 expressing S-6P from B.1.1.529/Omicron, a single intranasal dose of all three versions of B/HPIV3 vectors was protective against matched or heterologous WA1/2020, B.1.617.2/Delta or BA.1 (B.1.1.529.1)/Omicron challenge; hamsters were protected from challenge virus replication in the lungs, while low levels of challenge virus were detectable in the upper respiratory tract of a small number of animals. Immunization also protected against lung inflammatory response after challenge, with mild inflammatory cytokine induction associated with the slightly lower level of cross-protection of WA1/2020 and B.1.617.2/Delta variants against the BA.1/Omicron variant. Serum antibodies elicited by all vaccine candidates were broadly reactive against 20 antigenic variants, but the antigenic breadth of antibodies elicited by B/HPIV3-expressed S-6P from the ancestral or B.1.617.2/Delta variant exceeded that of the S-6P B.1.1.529/Omicron expressing vector. These results will guide development of intranasal B/HPIV3 vectors with S antigens matching circulating SARS-CoV-2 variants.

**Author Summary:** Intranasal COVID vaccines have the potential to stimulate respiratory mucosal immunity, effectively restricting replication of SARS-CoV-2 in the respiratory tract, thereby reducing virus shedding and transmission. To develop pediatric vaccines for intranasal immunization against HPIV3 and SARS-CoV-2, we use live-attenuated bovine-human parainfluenza virus vaccine, a pediatric intranasal parainfluenza virus vaccine candidate, designed to express the stabilized SARS-CoV-2 spike protein. We compared the immunogenicity and breadth of protection of the B/HPIV3-expressed ancestral, B.1.617.2/Delta, or B.1.1.529/Omicron variants following intranasal immunization in hamsters. All three B/HPIV3 vectors replicated in the respiratory tract, induced mucosal and serum anti-S IgA and IgG, and were protective in the hamster model against matched or heterologous WA1/2020, B.1.617.2/Delta or BA.1 (B.1.1.529.1)/Omicron SARS-CoV-2 challenge. Serum antibodies elicited by all intranasal vaccine candidates were broadly reactive against 20 antigenic variants of SARS-CoV-2, but the antigenic breadth of antibodies elicited by B/HPIV3-expressed stabilized S from the ancestral or B.1.617.2/Delta variant exceeded that of the S-6P B.1.1.529/Omicron expressing vector. Thus, these intranasal vectored SARS-CoV-2 vaccine candidates induce cross-protective SARS-CoV-2 immunity with antigenic breadth similar to that of injectable SARS-CoV-2 vaccines. These results will guide development of intranasal COVID vaccines based on B/HPIV3 vectors with S antigens matching current SARS-CoV-2 variants.

## Introduction

SARS-CoV-2 caused a pandemic with more than 775 million COVID-19 cases resulting at the time of writing in more than seven million deaths worldwide [1]. Since the emergence of SARS-CoV-2, children represent 17.9% of all cases in the United States (https://www.aap.org). While most infants and children generally are asymptomatic or exhibit mild to moderate symptoms, some may develop severe illness or complications. COVID-associated hospitalizations of children have substantially increased after emergence of Omicron variants [2, 3]. 90% of all hospitalized children were unvaccinated, indicating that lack of vaccination is the most important risk factor for severe disease in children [3, 4], similarly to adults. Vaccination and immunity from prior COVID infections mitigate risk of severe COVID in immunocompetent children [5].

mRNA-based vaccines are available for infants and young children six months of age and older. While these vaccines induce strong systemic immunity and are highly effective in preventing severe disease, they do not efficiently protect the upper respiratory tract from SARS-CoV-2 infection and their effectiveness against mucosal SARS-CoV-2 replication and transmission via the respiratory route is incomplete [6, 7]. Thus, intranasally delivered vaccines with the ability to directly stimulate respiratory mucosal immunity are needed that are effective in reducing breakthrough infections, and in restricting replication of SARS-CoV-2 in the respiratory tract, thereby reducing virus shedding and community transmission. Since mRNA vaccine-induced immunity wanes rapidly [8], intranasal vaccines that elicit mucosal immunity could be used alone or as heterologous boosters with mRNA-based vaccines to induce mucosal immunity, and to increase the antigenic breadth and duration of protection [9–12].

We previously developed vector vaccines for intranasal immunization based on chimeric bovine/human parainfluenza virus type 3 (B/HPIV3). B/HPIV3 had originally been developed as a live-attenuated pediatric vaccine for intranasal immunization against HPIV3 [13, 14], which is an important pediatric respiratory pathogen. B/HPIV3 contains the N, P, M and L genes of bovine PIV3, providing a strong host range restriction in humans. The HN and F proteins of B/HPIV3 are derived from human PIV3 and represent the major protective antigens of HPIV3. A B/HPIV3 vector expressing the fusion protein of respiratory syncytial virus (RSV) was previously evaluated as a bivalent pediatric vaccine candidate against both HPIV3 and RSV. B/HPIV3 and B/HPIV3 vectors expressing RSV F replicate to high titers in hamsters and in nonhuman primates (African green monkeys and rhesus macaques) [13, 15–19]. These vaccine candidates were highly restricted in replication in humans; in clinical studies, replication of B/HPIV3 vectors was undetectable or highly restricted in HPIV3 seropositive adults and children [14, 20]. In HPIV3-seronegative children 6-18 months of age, B/HPIV3 and B/HPIV3 expressing the RSV F protein were highly attenuated, and highly immunogenic against HPIV3 [14, 21], Clinicaltrials.gov NCT00686075].

Upon emergence of SARS-CoV-2, we evaluated B/HPIV3-vectored vaccines expressing the spike (S) protein of the ancestral SARS-CoV-2 strain stabilized in its prefusion form with two (S-2P, [22]) or six (S-6P, [23]) proline substitutions [24, 25]. B/HPIV3/S-6P was highly immunogenic and protective against the vaccine-matched ancestral SARS-CoV-2 as well as Alpha or Beta variants of concern in hamsters [25]. In rhesus macaques, intranasal/intratracheal administration of B/HPIV3/S-6P induced strong S-specific antibody and T cell responses systemically and in the airways, and was protective against a vaccine-matched SARS-CoV-2 challenge [26]. B/HPIV3/S-6P is currently being evaluated in a phase I clinical trial in adults (Clinicaltrials.gov NCT06026514), with plans for its sequential evaluation in HPIV3-seropositive and seronegative children and infants.

The continuous emergence of new SARS-CoV-2 variants requires updating of the S antigen expressed by the B/HPIV3 vector to optimize the immunogenicity of this vector vaccine. While antigenic breadth and cross-protection against different variants following immunization with injectable COVID vaccines has been characterized in many studies, less is known about the breadth of protection following intranasal immunization. In the present study, we generated B/HPIV3-vectored vaccines for intranasal immunization that express the S-6P versions of B.1.617.2/Delta or B.1.1.529/Omicron variants, and we explored their immunogenicity and the antigenic breadth of the systemic and mucosal antibody response in comparison to the previous version expressing the 6P stabilized S antigen of the ancestral SARS-CoV-2 strain. In addition, we evaluated their protective efficacy in hamsters against challenge with vaccine-matched or heterologous SARS-CoV-2 strains. The results will inform strategies to update the vectored S antigens to generate B/HPIV3-vectored clinical study material.

## Results

### Construction and rescue of B/HPIV3 expressing the S-6P version of the B.1.617.2/Delta and B.1.1.529/Omicron variants

Previously, we generated and evaluated in hamsters and rhesus macaques the B/HPIV3 vector vaccine candidate B/HPIV3/S-6P, expressing the SARS-CoV-2 S protein of the ancestral Wuhan-Hu-1 strain (GenBank MN908947), prefusion-stabilized by six proline substitutions [23, 25, 26]. In the present study, we generated B/HPIV3/S-Delta-6P and B/HPIV3/S-Omicron-6P expressing the full-length S proteins derived from SARS-CoV-2 B.1.617.2/Delta or B.1.1.529/Omicron variants, respectively, following the same strategy (Fig. 1A). The S-Delta-6P and S-Omicron-6P open reading frames (ORFs) (aa 1-1,273) were codon-optimized for human expression and include six proline substitutions to stabilize S in the prefusion form [22, 23]. Each S ORF was framed by nucleotide adapters containing the BPIV3 gene start and gene end signal sequences and placed as an additional gene between the N and P genes in the B/HPIV3 vector [25, 26] (Fig. 1A). B/HPIV3/S-Delta-6P and B/HPIV3/S-Omicron-6P were recovered from cDNA by reverse genetics and passaged once on Vero cells to generate passage 2 working stocks. Working stocks were titrated by dual-staining immunoplaque assay to confirm titers [6.2 and 6.3 log_10_ plaque-forming units (pfu) per ml for B/HPIV3/S-Delta-6P and B/HPIV3/S-Omicron-6P]. Ninety-five and 85% of the plaques generated by the B/HPIV3/S-Delta-6P and B/HPIV3/S-Omicron-6P stocks, respectively, were positive for HPIV3 and S antigen by dual-staining immunoplaque assay. Sanger sequencing of overlapping PCR fragments covering the entire genome (excluding regions of the genome ends complementary to the terminal sequencing primers) did not reveal any adventitious mutations.

**Fig 1.**
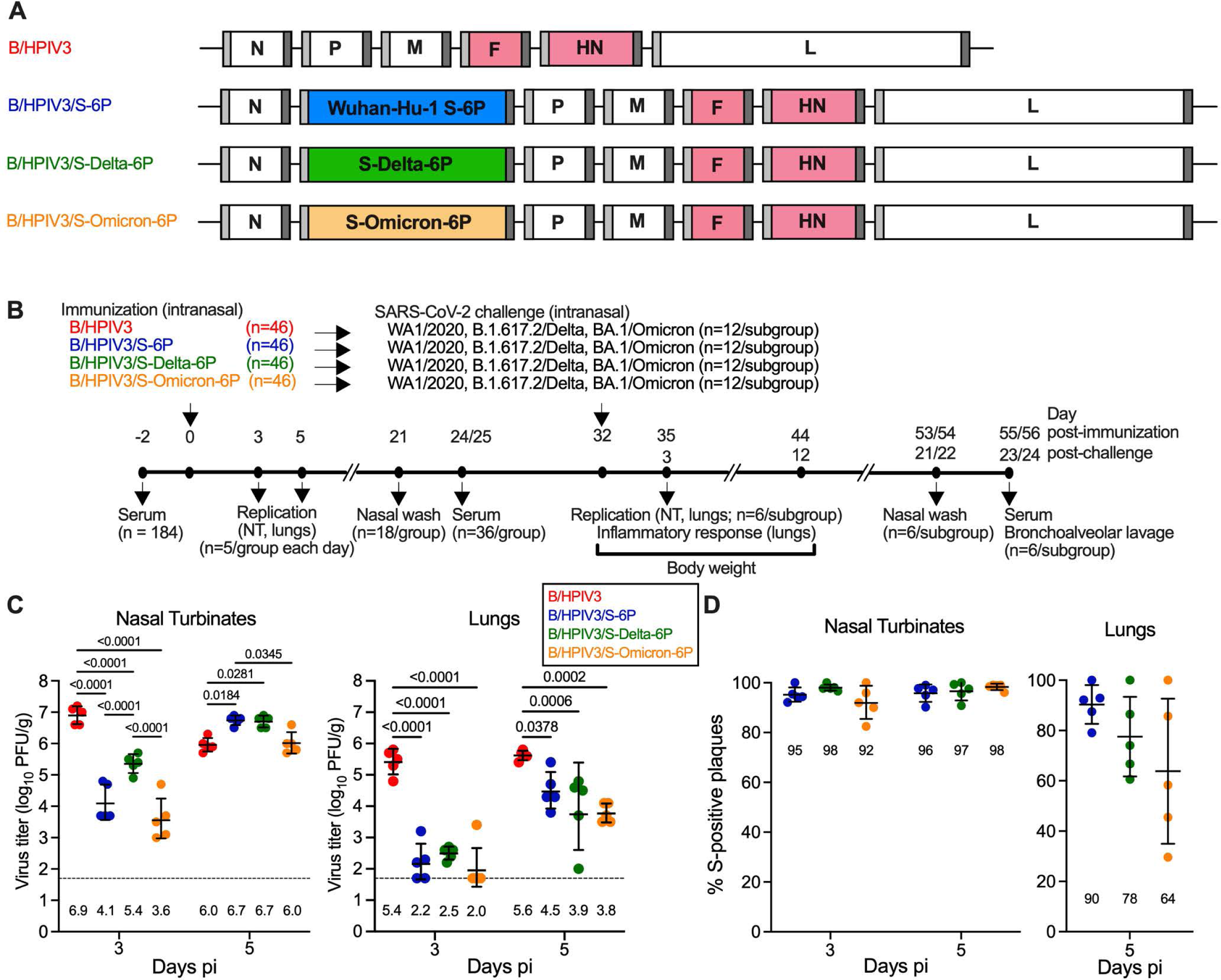
Replication of B/HPIV3, B/HPIV3/S-6P, B/HPIV3/S-Delta-6P and B/HPIV3/S-Omicron-6P in hamsters. (A) Genome map of the B/HPIV3 empty vector, B/HPIV3/S-6P expressing the S-6P stabilized version of the spike protein (S) of the ancestral SARS-CoV-2 strain (blue, [25, 26], and of the newly-generated versions B/HPIV3/S-Delta-6P (green) and B/HPIV3/S-Omicron-6P (orange). BPIV3 N, P, M and L genes are in white and HPIV3 F and HN genes in pink. Sequences corresponding to S ORFs (aa 1-1,273) from Wuhan-Hu-1, Delta or B.1.1.529 strains were codon-optimized for expression in humans, the furin cleavage site “RRAR” was removed and replaced by “GSAS” [22], and six proline substitutions were introduced to stabilize the S antigen in the prefusion form [23] (not shown). An additional gene encoding the respective S ORF, flanked by a BPIV3 gene start (light grey) and gene end (dark grey) signal sequence, was inserted between the BPIV3 N and P genes [25, 26]. (B) Timeline of the hamster experiment. Five to six-week-old hamsters in groups of 46 were intranasally immunized with 5 log_10_ pfu of B/HPIV3 empty vector or B/HPIV3 expressing the S-6P, S-Delta-6P or S-Omicron-6P. (C) On days 3 and 5 post-immunization (pi), five hamsters per group each day were euthanized, and vaccine titers were determined from nasal turbinates (C, left panel) and lungs (C, right panel) by dual-staining immunoplaque assay. The limit of detection (dotted line) is 50 pfu/g of tissue. Geometric mean titers (GMTs) for each group are indicated above x axes. Two-way ANOVA with Sidak post-test; exact p values are indicated for levels of significance p<0.05. (D) Percentages of S-positive plaques in nasal turbinates (left panel) and lungs (right panel) on days 3 and 5 pi for nasal turbinates and day 5 pi only for lungs. Due to low level of replication on day 3 pi, lung samples were not investigated. Each animal is represented by a circle and GMTs with geometric standard deviations are shown.

### B/HPIV3/S-Delta-6P and B/HPIV3/S-Omicron-6P replicate efficiently in the respiratory tract of hamsters

The replication, immunogenicity, and protective efficacy of B/HPIV3/S-Delta-6P and B/HPIV3/S-Omicron-6P were evaluated in five- to six-week-old golden Syrian hamsters, and B/HPIV3/S-6P and the B/HPIV3 empty vector were included as controls (see Fig. 1B for timeline of the experiment). On day 0, four groups of 46 hamsters each were immunized intranasally with 5 log_10_ pfu of B/HPIV3, B/HPIV3/S-6P, B/HPIV3/S-Delta-6P, or B/HPIV3/S-Omicron-6P. On days 3 and 5 post-immunization (pi), five hamsters per group were euthanized and nasal turbinates (NTs) and lungs were harvested. Tissue homogenates were prepared, and vaccine virus titers were determined by immunoplaque assay.

Similarly to previous studies, the B/HPIV3 empty vector replicated to high titers on day 3 pi (geometric mean titers [GMTs] of 6.9 and 5.4 log_10_ pfu/g in NTs and lungs, respectively, Fig. 1C, left and right panel). On day 5 pi, B/HPIV3 titers were slightly lower than on day 3 in NTs (6.0 log_10_ pfu/g), but remained at a level similar to day 3 in lungs (5.6 log_10_ pfu/g). On day 3, the GMTs of all S-expressing versions (B/HPIV3/S-6P, B/HPIV3/S-Delta-6P, and B/HPIV3/S-Omicron-6P) were significantly lower than those of the empty B/HPIV3 vector [between 32- and 1,995-fold (NTs) and 794- and 2,512-fold (lungs), p <0.0001]. By day 5 pi, titers in NTs of the three S-expressing vectors had increased to levels comparable to those in B/HPIV3-infected hamsters on day 3 (Fig. 1C, left panel). In the lungs, titers of all S-expressing B/HPIV3 vectors had also increased by day 5 (GMTs ranging from 3.8 to 4.5 log_10_ pfu/g), but peak titers remained lower than those of B/HPIV3 (13-fold for B/HPIV3/S-6P; 79-fold for B/HPIV3-S-Delta-6P and B/HPIV3/S-Omicron-6P). Thus, replication of B/HPIV3-S expressing vectors was delayed in the NTs and delayed and reduced in the lungs due to the insertion of the foreign S gene into the virus genome.

The stability of the S expression by the B/HPIV3 vectors in NTs and lungs was also evaluated using a dual staining immuno-plaque assay (Fig. 1D left and right panel). Between 92 and 98% of the plaques from NT-derived B/HPIV3/S-Delta-6P, B/HPIV3/S-Omicron-6P or B/HPIV3/S-6P expressed S. This suggests substantial stability of S expression in the upper airways (Fig. 1D, left panel). In lung homogenates, between 64 and 90% of the plaques generated by the B/HPIV3 vectors expressed S, suggesting more variable S expression among the different vectors in the lower airways, and greatest variability of S expression by B/HPIV3/S-Omicron-6P (Fig. 1D, right panel). Due to the low titers of B/HPIV3 S-expressing vectors in the lungs on day 3 pi, stability of S expression on day 3 in lungs was not evaluated.

### B/HPIV3/S-Delta-6P and B/HPIV3/S-Omicron-6P induced mucosal and serum anti-S antibody responses in hamsters

We next evaluated the mucosal antibody responses induced by the B/HPIV3 S-expressing vectors using nasal washes collected on day 21 pi from 18 of the 36 remaining hamsters per immunized group that were selected randomly (Fig. 2A). S-specific IgG and IgA titers were determined by IgG/IgA ELISA assay based on S-6P antigen derived from the ancestral Wuhan-Hu-1 strain. This dual ELISA allows detection of IgA and IgG in the same sample by sequential reads (see materials and methods). Using the ancestral S antigen, we detected mucosal anti-S IgG in nasal washes from 8/18 and 7/18 hamsters immunized with B/HPIV3/S-6P or B/HPIV3/S-Delta-6P, respectively (GMT ELISA titer of 1.3 and 1.2 log_10_, respectively). No hamsters immunized with B/HPIV3/S-Omicron-6P had IgG antibodies to the ancestral version of S detectable in nasal washes (Fig. 2A, left panel). However, 17/18, 12/18 and 4/18 hamsters immunized with B/HPIV3/S-6P, B/HPIV3/S-Delta-6P or B/HPIV3/S-Omicron-6P, respectively, had mucosal anti-S IgA detectable in nasal washes (GMT ELISA titer from 1.04 to 1.7 log_10_), indicating that all S-expressing vectors induced a mucosal anti-S IgA response (Fig. 2A, right panel), even though B/HPIV3/S-Omicron-6P was significantly less efficient in inducing a mucosal anti-S antibody response to the ancestral version of S, with a high animal-to-animal variability. As expected, no mucosal anti-S IgG or IgA antibody response was detectable in hamsters immunized with the B/HPIV3 empty-vector control.

**Fig 2.**
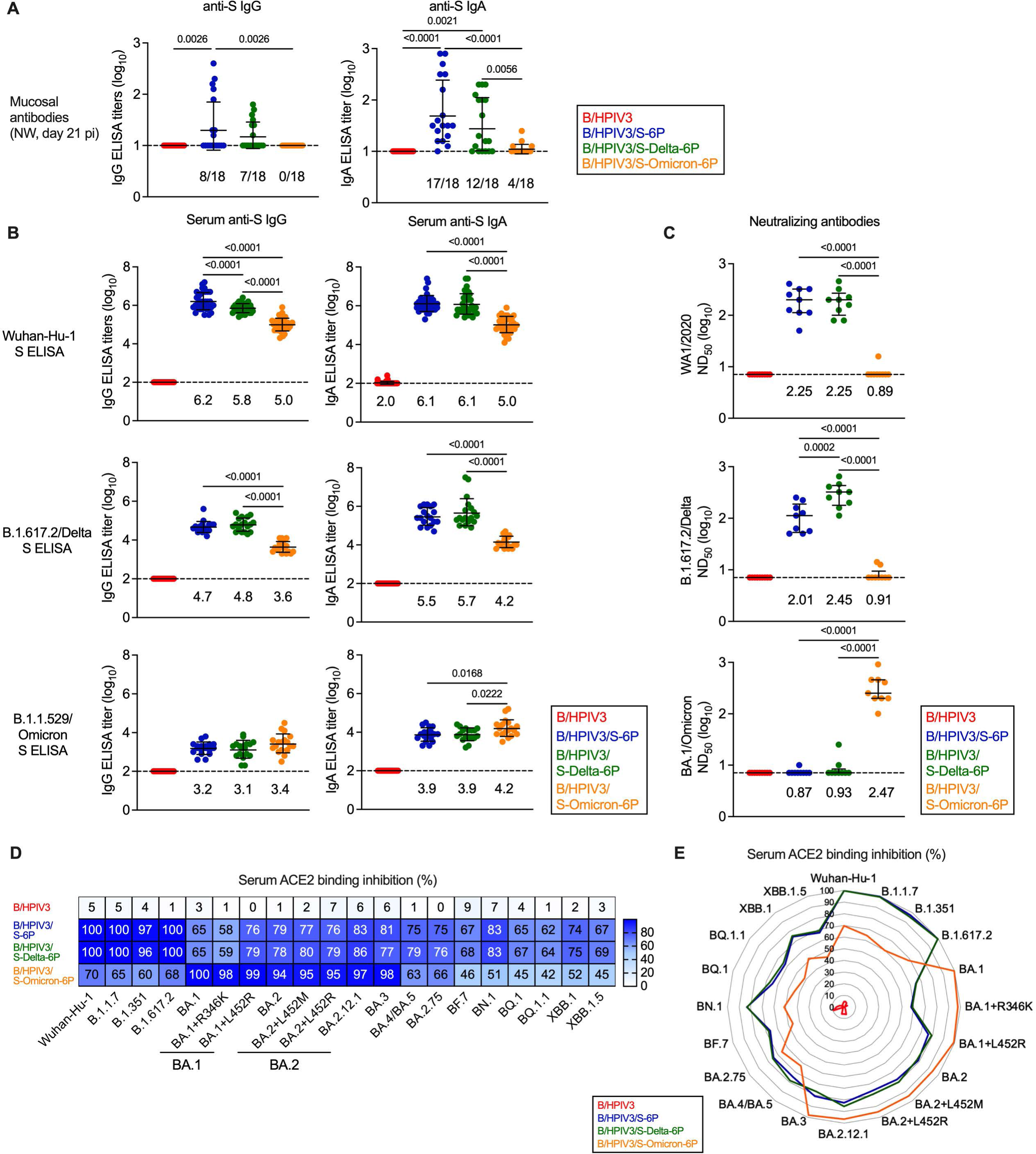
Mucosal and serum antibody responses in immunized hamsters. (A) IgG (left panel) and IgA (right panel) anti-S mucosal antibody titers were evaluated by ELISA from nasal washes collected on day 21 post-immunization (pi, n = 18 hamsters per group picked at random). The fraction of animals with titers above the limit of detection is indicated above x axes. (B-E) On day 24 or 25 pi, serum was collected from n = 36 hamsters per group. (B) The IgG (left panels) and IgA (right panels) anti-S antibody titers were evaluated by ELISA using purified preparations of S antigen from the Wuhan-Hu-1 strain (top row), B.1.617.2/Delta (middle row) or B.1.1.529/Omicron variants (bottom row). ELISA results using purified RBD antigen preparations from the Wuhan-Hu-1 strain are shown in Fig. S1. The limit of detection of reciprocal antibody titers (dotted line) is 1 log_10_ (A) or 2 (B) log_10_. (C) Sera collected on day 24 or 25 pi were also analyzed for SARS-CoV-2 neutralizing antibody titers. Live virus neutralization assays were performed using the WA1/2020 strain or B.1.617.2/Delta or BA.1/Omicron variants. Titers were expressed as 50% neutralizing doses. (A-C) One-way ANOVA with Tukey or Sidak post-test; exact p values are indicated for levels of significance p<0.05. Each hamster is represented by a symbol and medians with interquartile ranges are shown. (B-C) GMTs are indicated above x axes. (D-E) Binding inhibition of soluble ACE2 protein to SARS-CoV-2 S proteins from 20 different variants by serum antibodies from immunized hamsters. Data are represented as a heatmap with the median percent inhibition (from n = 36 hamsters per group) of ACE2 binding relative to a non-serum control indicated for each variant, (D) or as a radar plot, with each segment representing a variant, and percent inhibition indicated by concentric circles (E).

We also determined the serum IgG and IgA titers elicited by the B/HPIV3 vectors to the S proteins of the ancestral strain as well as B.1.617.2/Delta and B.1.1.529/Omicron variants at day 24 or 25 pi using dual IgG/IgA ELISAs. Purified S protein from the indicated strains were used as antigens (Fig. 2B). An additional ELISA was performed to evaluate antibodies binding the receptor binding domain (RBD, derived from the ancestral strain; Fig. S1).

All three S-expressing B/HPIV3 vectors elicited robust serum IgG and IgA responses to both Wuhan-Hu-1 S and RBD (IgG and IgA ELISA GMT between 5.0-6.2 log_10_, and 3.5-5.8 for S and RBD, respectively, Fig. 2B top panels and Fig. S1). However, serum IgG and IgA GMTs to the ancestral version of the S protein elicited by B/HPIV3/S-Omicron-6P were 6- to 16-fold lower than those elicited by the matched vaccine B/HPIV3/S-6P- or by B/HPIV3/S-Delta-6P (p<0.0001, Fig. 2B, top panels), and anti-RBD IgG and IgA GMTs elicited by B/HPIV3/S-Omicron-6P were 31 to 88-fold lower (p<0.0001, Fig. S1), reflecting antigenic differences between S and RBD proteins of the Wuhan-Hu-1, Delta, and B.1.1.529/Omicron variants [27].

The homologous and heterologous antibody responses induced by each B/HPIV3-vectored S antigen to the S protein from Delta or B.1.1.529/Omicron variants were also determined (Fig. 2B, middle and bottom panels), showing that B/HPIV3/S-6P and B/HPIV3/S-Delta-6P-elicited antibodies bound significantly stronger to the Delta S protein than those elicited by B/HPIV3/S-Omicron-6P. Overall lower IgG titers were detected when S-2P of the B.1.1.529/Omicron variant was used as the antigen; B.1.1.529/Omicron anti-S IgG GMTs were comparable in all groups, while B.1.1.529/Omicron anti-S IgA GMTs elicited by B/HPIV3/S-6P and B/HPIV3/S-Delta-6P were significantly lower than those in B/HPIV3/S-Omicron-6P immunized hamsters (Fig. 2B, bottom panel). As expected, no anti-S or anti-RBD IgG/IgA antibodies were induced in hamsters immunized by the B/HPIV3 empty-vector control (Fig. 2B, S1).

The 50% serum neutralizing antibody titers (ND_50_) to SARS-CoV-2 variants corresponding to the three different B/HPIV3-expressed S antigens (WA1/2020, B.1.617.2/Delta, and B.1.1.529/Omicron) were determined at BSL3 by ND_50_ assay for nine randomly selected hamsters from each group (Fig. 2C). As expected, B/HPIV3/S-6P induced strong WA1/2020-neutralizing titers, similarly to B/HPIV3/S-Delta-6P (GMTs of 2.25 log_10_ in both groups, Fig. 2C, top panel), while only one of the nine B/HPIV3/S-Omicron-6P immunized hamsters had WA1/2020-neutralizing antibodies detectable. Against SARS-CoV-2 of the B1.617.2/Delta variant, the matched vaccine candidate B/HPIV3/S-Delta-6P induced significantly higher neutralizing titers than B/HPIV3/S-6P (GMT 2.45 vs 2.01, P=0.0002), while only two of nine B/HPIV3/S-Omicron-6P immunized animals had very low B.1.617.2/Delta neutralizing serum antibody titers detectable. However, using SARS-CoV-2 BA.1/Omicron, exclusively the sera from B/HPIV3/S-Omicron-6P immunized hamsters had robust neutralizing antibody titers detectable, while only a single B/HPIV3/S-6P- and two B/HPIV3/S-Delta-6P-immunized hamsters had very low BA.1/Omicron ND_50_ titers detectable. Thus, each vector induced strong neutralizing serum antibodies against its best-matching SARS-CoV-2 variant. There was substantial cross-neutralization of antibodies elicited by the ancestral and the Delta antigens between the ancestral strain and Delta variant, but this was not true for the BA.1/Omicron variant. No SARS-CoV-2 neutralizing antibodies were detected in sera from B/HPIV3 control immunized hamsters.

To evaluate further the antigenic breadth of the vaccine candidates, we evaluated the sera in an ACE2 binding inhibition assay as a surrogate to live virus neutralization assays. This assay evaluates the ability of serum antibodies to inhibit binding of soluble ACE2 receptor to S proteins derived from 20 different SARS-CoV-2 variants, including the recently circulating XBB variants (Fig. 2D and E). Results from this assay are expressed as % inhibition of ACE2 binding relative to a non-serum buffer-only negative control and are shown as a heat map (Fig. 2D, n = 36 per immunized group). The assay allows evaluation of the breadth of the serum antibody responses using relatively small volumes of serum. We found that sera from B/HPIV3/S-6P- and B/HPIV3/S-Delta-6P immunized hamsters highly efficiently inhibited ACE2 binding to S from the vaccine-matched strains (Wuhan-Hu-1 and B.1.617.2/Delta), but also of early variants, B.1.1.7/Alpha and B.1.351/Beta (96 to 100 median % inhibition; Fig. 2D and E). Interestingly, the binding inhibition of S proteins from BA.1 and BA.1+R346K variants by these sera was moderate (58 to 65 median % inhibition), while their ability to inhibit binding of ACE2 to S from BA.1+L452R, BA.2, BA.3, BA.4, BA.5, BQ.1, BF.7, BN.1 variants and derivatives as well as the recently circulating XBB variants was stronger (62 to 86% median inhibition). The results for each vaccine candidate were combined in a radar plot (Fig. 2E), showing that the breadth of serum antibodies elicited by B/HPIV3/S-6P and B/HPIV3/S-Delta-6P was comparable, with strongest reactivity to variants of the B.1.1.7/Alpha, B.1.351/Beta, and B.1.617.2/Delta lineages.

In contrast, serum antibodies in hamsters immunized with B/HPIV3/S-Omicron-6P showed only moderate inhibition against these pre-Omicron strains (60 to 70 median % inhibition, Fig. 2D and E), but they efficiently inhibited ACE2 binding of S proteins from the BA.1, BA.2, BA.3 Omicron sublineages and derivatives (94 to 100 median % inhibition).

However, their ACE2 binding inhibition of S proteins from more recent SARS-CoV-2 variants (BA.4/5, BA.2.75, BF7, BN.1, BQ.1, XBB.1 and derivatives) was lower than that of hamsters immunized with B/HPIV3/S-6P and B/HPIV3/S-Delta-6P, with median % inhibition ranging between 42% and 66%. As expected, sera from B/HPIV3 empty vector immunized hamsters did not inhibit binding of ACE2 to S from the 20 different variants analyzed.

### Weight change and lung inflammatory response of immunized hamsters following challenge with SARS-CoV-2 WA1/2020, BA1.617.2/Delta, or BA.1/Omicron variants

We next evaluated the protective efficacy of the B/HPIV3-S expressing vectors against challenge with homologous or heterologous SARS-CoV-2 variants (see Fig. 1B for timeline). To do so, the remaining 36 hamsters from each immunization group (B/HPIV3 empty-vector control, B/HPIV3/S-6P, B/HPIV3/S-Delta-6P, or B/HPIV3/S-Omicron-6P) were transferred to a BSL3 facility. Hamsters from each group were randomly distributed into three subgroups (n=12 per subgroup), and, on day 32 pi, challenged with 4.5 log_10_ 50% tissue culture infectious doses (TCID_50_) of SARS-CoV-2 of either the WA1/2020 or B.1.617.2/Delta variants, or the BA.1/Omicron variant, representing the major circulating variant at the time the study was designed.

After SARS-CoV-2 challenge, hamsters were weighed daily for 12 days to monitor for weight loss (Fig. 3A). Animals immunized with the B/HPIV3 empty-vector control and challenged with WA1/2020 or B.1.617.2/Delta exhibited moderate weight loss from day 2 to 8 post-challenge (pc; 14 and 7 median % weight loss on day 8, respectively) before re-gaining weight from day 8 to 12 pc (Fig. 3A, left and middle panels). Animals immunized with the S-expressing B/HPIV3 vectors were all protected from weight loss following challenge with WA1/2020 (p<0.001 from day 2 to 7 pc, p<0.01 on day 8 pc and p<0.01 on day 11 and 12 pc compared to the B/HPIV3 empty vector) or B.1.617.2/Delta (p<0.001 on day 2 and 3 pc compared to the B/HPIV3 empty vector) (Fig. 3A, left and middle panels). Following challenge with BA.1/Omicron, no weight loss was observed in the empty-vector control group, nor in any hamsters immunized with an S-expressing B/HPIV3 vector (Fig. 3A, right panel). This was expected, as previous studies have shown that BA.1/Omicron does not induce weight loss in hamsters [28].

**Fig 3.**
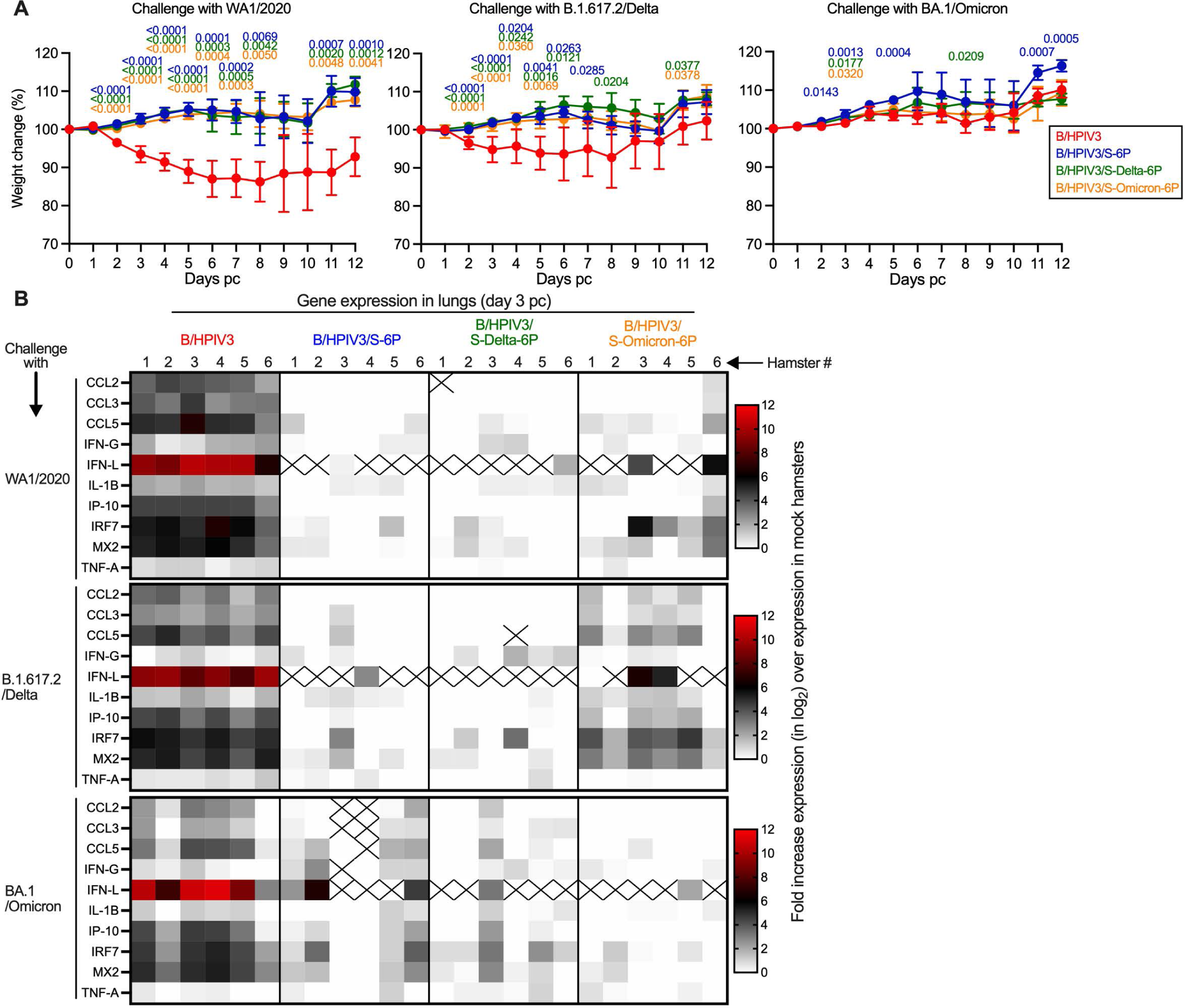
Weight change and lung inflammatory responses of immunized hamsters upon SARS-CoV-2 challenge. On day 32 pi, the 36 remaining hamsters per immunized group were sub-divided into three subgroups of 12 hamsters each. Each subgroup was challenged with 4.5 log_10_ TCID_50_ per animal of SARS-CoV-2 WA1/2020, B.1.617.2/Delta or BA.1/Omicron (see Fig. 1B for timeline). (A) Body weights were monitored daily from day 0 to 12 post-challenge (pc). Data are shown as mean percent body weight relative to the day 0 weight (n=12 hamsters/subgroup from day 0 to 3, n=6 hamsters/subgroup from day 4 to 12 except for B/HPIV3-immunized and WA1-challenged subgroup with 4 or 5 animals from day 8 to 12). Mixed-effects analysis with Dunnett post-test; exact p values are indicated for levels of significance p<0.05. (B) Expression of inflammatory/antiviral genes in lungs on day 3 pc. On day 3 pc, six hamsters per subgroup were euthanized and lungs were harvested and homogenized. Total RNA was extracted from lung homogenates and the expression of 10 inflammatory/antiviral genes was evaluated by RT-qPCR using TaqMan assays [25]. Results were analyzed using ΔΔCt method and normalized to beta actin. Relative expression was expressed as log_2_ fold change over the expression level in five unimmunized, unchallenged hamsters [25] and represented as heatmaps with one heatmap per challenge virus. Non-detected genes are crossed.

Six hamsters per subgroup were euthanized on day 3 pc, and lungs and nasal turbinates were collected. To assess effects of SARS-CoV-2 challenge further, we determined lung inflammatory cytokine responses by measuring the expression of 10 inflammatory/antiviral genes by RT-qPCR in their lungs. Data represented as heatmaps show the fold increase of gene expression (in log_2_) over the mean expression in five mock-immunized, mock-challenged hamsters that were derived from a separate previous study [25] (Fig. 3B). Upon challenge with WA1/2020, B.1.617.2/Delta or BA.1/Omicron, a strong increase in expression of inflammatory/antiviral genes was detected in the lungs of B/HPIV3-immunized hamsters (1 to 12 log_2_ fold-increase in expression, with CCL5, IFN-L, IRF7 and MX2 being the most-upregulated genes) (Fig. 3B, B/HPIV3 panels).

No or low increases in expression of inflammatory/antiviral genes were detected in B/HPIV3/S-6P- or B/HPIV3/S-Delta-6P-immunized hamsters after challenge with either homologous or heterologous SARS-CoV-2 strains WA1/2020 or B.1.617.2/Delta. However, following heterologous challenge of these B/HPIV3/S-6P- or B/HPIV3/S-Delta-6P immunized animals with the more antigenically distant BA.1/Omicron variant, moderate expression of some inflammatory/antiviral genes were observed in 4/6 and 2/6 hamsters, respectively (Fig. 3B, B/HPIV3/S-6P and B/HPIV3/S-Delta-6P columns, bottom row). This suggested that immunization with B/HPIV3/S-6P or B/HPIV3/S-Delta-6P was less effective against lung inflammatory responses after BA.1/Omicron challenge.

On the other hand, no or low increases in expression of inflammatory/antiviral genes were detected in B/HPIV3/S-Omicron-6P immunized hamsters after homologous BA.1/Omicron challenge (Fig. 3B, B/HPIV3/S-Omicron-6P column, bottom row), while a moderate to strong increase in expression of inflammatory/antiviral genes was detected in the lungs of B/HPIV3/S-Omicron-6P-immunized hamsters following heterologous WA1/2020 or B.1.617.2/Delta virus challenge (Fig. 3B, B/HPIV3/S-Omicron-6P column, top and middle rows), suggesting that immunization with B/HPIV3/S-Omicron-6P was less effective against challenge with the heterologous WA1/2020 or B.1.617.2/Delta variant. These results are consistent with the greater antigenic distance between BA.1/Omicron and the ancestral WA1/2020 strain or the B.1.617.2/Delta variant.

### Replication of SARS-CoV-2 WA1/2020, B.1.617.2/Delta or BA.1/Omicron challenge virus in immunized hamsters

We also evaluated SARS-CoV-2 challenge virus replication in nasal turbinates (NTs) and lungs from these six hamsters per challenge subgroup that had been euthanized on day 3 pc. Challenge viral loads were first determined from lung tissue homogenates by RT-qPCR using previously-described TaqMan assays targeting SARS-CoV-2 subgenomic E (sgE) or N (sgN) mRNA as indicators for active challenge virus replication. As expected, WA1/2020 and B.1.617.2/Delta challenge virus replicated efficiently in the lungs of hamsters previously immunized with the B/HPIV3 empty vector (GMTs from 8.6 to 10.2 log_10_ copies of sgE or sgN/g, respectively, of lung tissues) (Fig. 4A, left and middle panels). BA.1/Omicron also replicated efficiently in the empty-vector control immunized hamsters, albeit to a lower level (GMT of 6.9 and 8.8 log_10_ copies of sgE and sgN/g of tissues, respectively) (Fig. 4A, right panel).

**Fig 4.**
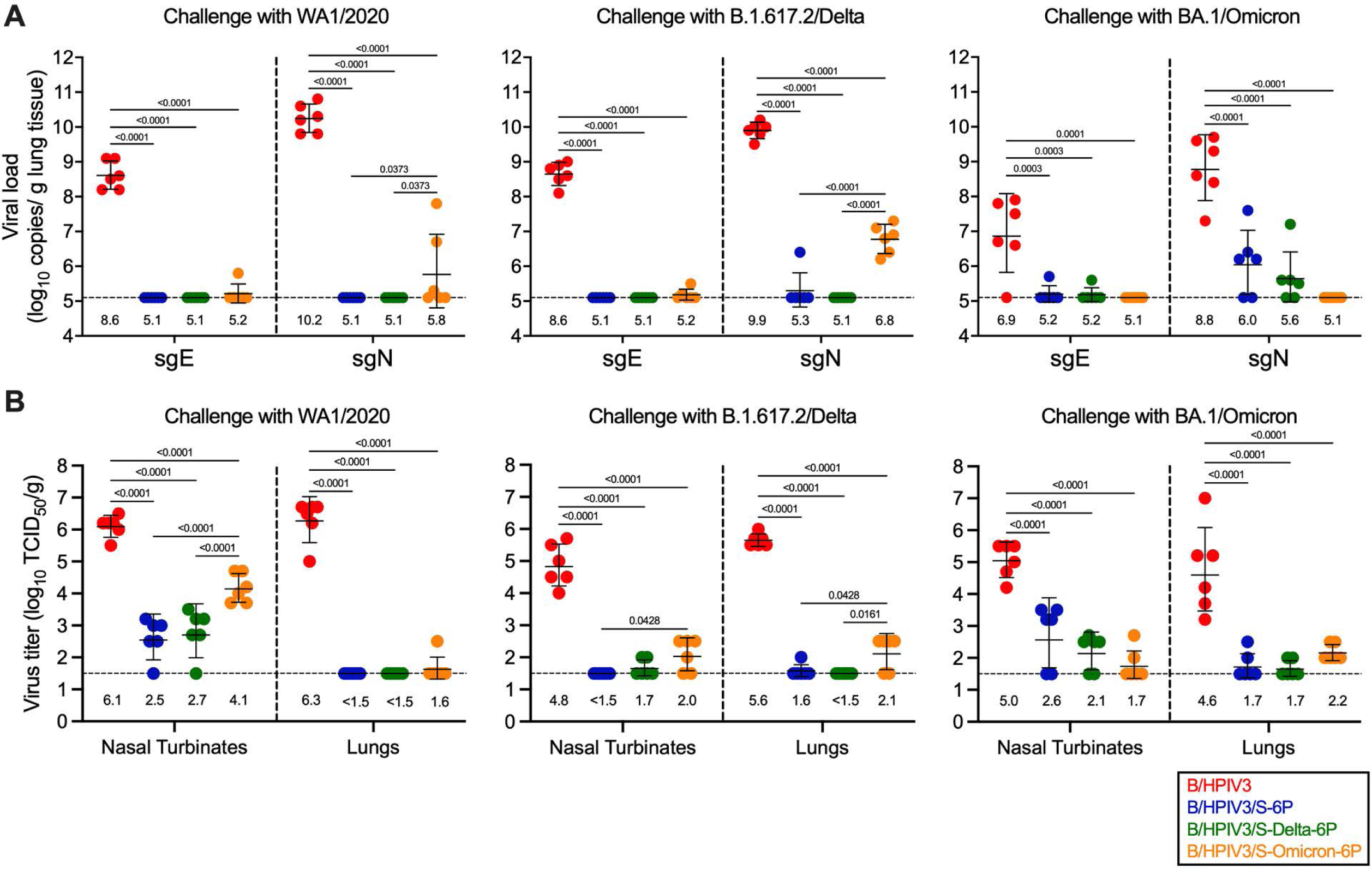
Protection of immunized hamsters from SARS-CoV-2 challenge virus replication. On day 3 pc, 6 of 12 hamsters per subgroup were euthanized and nasal turbinates and lungs were harvested, and homogenized. Aliquots were used to evaluate SARS-CoV-2 challenge virus replication by RT-qPCR (A) or viral culture (B). (A) Total RNA was extracted from aliquots of lung homogenates and SARS-CoV-2 viral loads were determined by RT-qPCR using TaqMan assays to quantify subgenomic E or N mRNA (Limit of detection: 5.1 log_10_ copies per g, dotted line). (B) Titers of SARS-CoV-2 challenge viruses, WA1/2020, B.1.617.2/Delta or BA.1/Omicron were determined from aliquots of nasal turbinate and lung homogenates and expressed in log_10_ TCID_50_ per g of tissue (Limit of detection: 1.5 log_10_ TCID_50_ per g, dotted line). Each hamster is represented by a symbol and GMTs with standard deviations are shown. GMTs are also indicated above x axes. One-way ANOVA with Tukey or Sidak post-test; exact p values are indicated for levels of significance p<0.05.

After challenge with WA1/2020 or B.1.617.2/Delta, hamsters immunized with B/HPIV3/S-6P or B/HPIV3/S-Delta-6P had no detectable virus replication in the lungs, indicating complete protection, except for one hamster immunized with B/HPIV3/S-6P that exhibited a low level of B.1.617.2/Delta sgN RNA (Fig. 4A, middle panel). However, in 2/6 B/HPIV3/S-Omicron-6P-immunized hamsters challenged with WA1/2020, sgE/N RNA was detected, albeit at a lower level than in empty-vector immunized animals (GMT of 5.8 log_10_ vs 10.2 log_10_ copies/g of sgN, Fig. 4A, left panel). In addition, all six B/HPIV3/S-Omicron-6P-immunized hamsters in the B.1.617.2/Delta challenge subgroup had sgN RNA detectable in the lungs (GMT of 6.8 log_10_ copies/g of sgN), albeit greatly reduced compared to the B.1.617.2/Delta viral load in empty-vector immunized hamsters (Fig. 4A, middle panel). This indicates that that immunization with B/HPIV3/S-Omicron-6P conferred less-than-complete protection against WA1/2020 or B.1.617.2/Delta challenge virus replication in the lungs, reflecting the antigenic distance between BA.1/Omicron and WA1/2020 or B.1.617.2/Delta variants.

In BA.1/Omicron-challenged subgroups of B/HPIV3/S-6P and B/HPIV3/S-Delta-6P immunized hamsters, sgE or sgN was detected in 4/6 immunized hamsters per subgroup, albeit at a low level (GMT of 6.0 and 5.6 log_10_ copies/g of sgN, for B/HPIV3/S-6P and B/HPIV3/S-Delta-6P immunized hamsters, respectively), indicating less-than-complete protection against heterologous BA.1/Omicron challenge. On the other hand, animals in BA.1/Omicron-challenged subgroups immunized with the matched vector B/HPIV3/S-Omicron-6P had no sgE or sgN detectable in lungs and appeared fully protected from challenge virus replication (Fig. 4A, right panel).

We also evaluated SARS-CoV-2 replication in NT and lungs on day 3 pc by titration of clarified tissue homogenates on Vero cells. As expected, in NT and lung homogenates from B/HPIV3 empty-vector immunized animals, WA1/2020, B.1.617.2/Delta and BA.1/Omicron challenge virus was detectable at substantial titers (between 4.8-6.1 log_10_ and 4.6-6.3 log_10_ TCID_50_/g in NT and lungs, respectively) (Fig. 4B), indicating that these variants replicated efficiently in B/HPIV3 empty vector immunized hamsters.

Following homologous or heterologous challenge with WA1/2020 or B.1.617.2/Delta variants (Fig. 4B, left and middle panels), hamsters from the B/HPIV3/S-6P- and B/HPIV3/S-Delta-6P-immunized groups had no or relatively low levels of challenge virus detectable in the NTs (undetectable to 2,500-fold reduced titers compared to hamsters in B/HPIV3 empty-vector control group, p<0.0001); in their lungs, challenge virus was undetectable, with the exception of one hamster in the B/HPIV3/S-6P immunized group challenged with B.1.617.2/Delta (p<0.0001 compared to B/HPIV3 empty vector, Fig. 4B, left and middle panels). Following BA.1/Omicron challenge (Fig. 4B, right panel), virus titers in NT of the B/HPIV3/S-6P- and B/HPIV3/S-Delta-6P-immunized hamsters were 250-fold lower than in the B/HPIV3 empty-vector immunized group (Fig. 4B, right panel, p<0.0001). In lungs collected from animals in these two immunization groups, BA.1/Omicron was detectable only in two hamsters per group at low levels close to the limit of detection (Fig. 4B, right panel, p<0.0001 compared to B/HPIV3 empty vector). Thus, hamsters immunized with B/HPIV3/S-6P- and B/HPIV3/S-Delta-6P were robustly but nevertheless less effectively protected against BA.1/Omicron than against WA1/2020 or B.1.617.2/Delta.

On the other hand, all hamsters immunized with B/HPIV3/S-Omicron-6P and challenged with the ancestral WA1/2020 had challenge virus detectable in NT (GMT 100-fold reduced compared to B/HPIV3 empty-vector immunized hamsters, p<0.0001, Fig. 4B, left panel). However, in their lungs, with the exception of one hamster that had low titers of WA1/2020 detectable, no WA1/2020 virus was detectable, indicating substantial protection by B/HPIV3/S-Omicron-6P against the ancestral WA1/2020. In addition, low levels of B.1.617.2/Delta and BA.1/Omicron challenge virus were detected in the NT and lungs of the B/HPIV3/S-Omicron-6P-immunized hamsters (Fig. 4B, middle and right panels).

### Magnitude and breadth of anamnestic mucosal and serum antibody responses upon SARS-CoV-2 challenge

In the remaining six hamsters per challenge subgroup, we evaluated mucosal and serum anamnestic immune responses following challenge. To do so, nasal washes were collected on day 21/22 pc from the same hamsters that had NW collected on day 21 pi. On day 23/24 pc, animals were necropsied and bronchoalveolar lavage and sera were collected (see Fig. 1B for timeline). Anti-S and anti-RBD IgG and IgA titers were then determined by ELISA using S antigen from the Wuhan-Hu-1 strain (Fig. 5A-C, Fig. S2). The breadth of the anamnestic serum antibody response was evaluated further in an assay that measures binding inhibition of soluble ACE2 receptor to S antigens from 20 variants (Fig. 5D and 5E).

**Fig 5.**
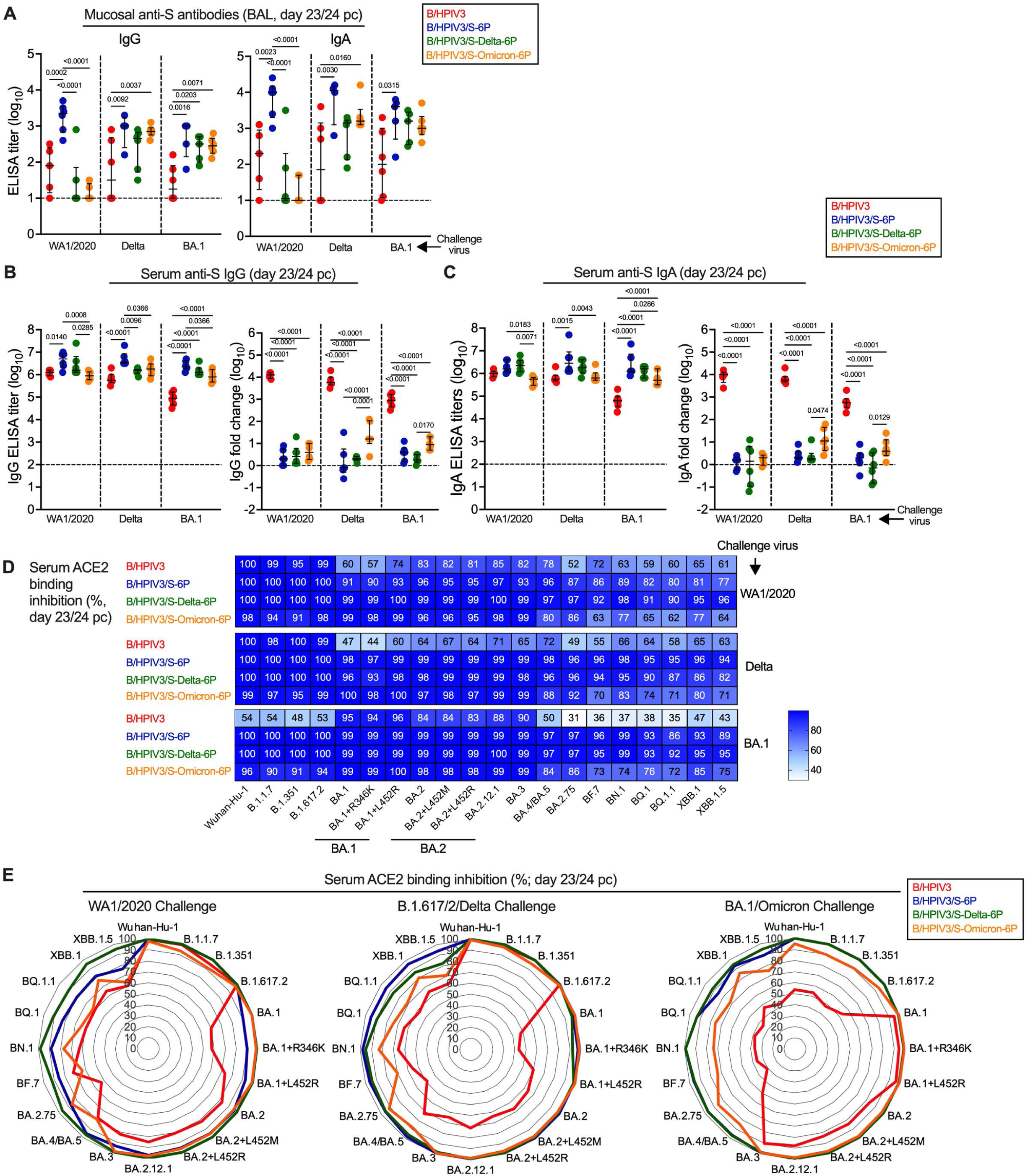
Mucosal and serum antibody responses in immunized hamsters upon SARS-CoV-2 challenge. On day 23 or 24 pc, the six remaining hamsters per subgroup were euthanized, and bronchoalveolar lavage (BAL) and sera were collected (see Fig. 1B for timeline of the experiment) for evaluation of the antibody response by ELISA (A-C) or ACE2 binding inhibition assay (D-E). (A) Anti-S IgG (left panel) and IgA (right panel) antibody titers in BAL (limit of detection: 1 log_10_, dotted line). N = 6 hamsters per subgroup with the exception of n = 5 for B/HPIV3 immunized/WA1/2020 challenged, B/HPIV3/S-6P-immunized/BA.1 challenged, B/HPIV3/S-Delta-6P-immunized/BA.1 challenged, B/HPIV3/S-Omicron-6P-immunized/WA1/2020 challenged and n = 4 for B/HPIV3/S-6P-immunized/Delta challenged. (B-C) Serum anti-S IgG (B) and IgA (C) titers after challenge (left panels) and fold changes (in log_10_) of post-challenge titers over post-immunization titers (right panels). N= 6 with the exception of n = 5 for B/HPIV3-immunized/WA1/2020 challenged. Purified preparations of S from the Wuhan-Hu-1 strain were used as antigens in the ELISA assays. Each hamster is represented by a symbol and medians with interquartile ranges are shown. One-way ANOVA with Tukey post-test; exact p values are indicated for levels of significance p<0.05. (D-E) Binding inhibition of soluble ACE2 protein to SARS-CoV-2 S proteins from 20 different variants by serum antibodies from immunized and challenged hamsters. Data are represented as a heatmap with one heatmap per challenge virus and with the median percent inhibition of ACE2 binding relative to a non-serum control indicated (D) or as radar plots (E). N= 6 with the exception of n = 5 for B/HPIV3-immunized/WA1/2020 challenged.

Using the ancestral Wuhan-Hu-1 S antigen, we detected only low to background levels of nasal anti-S IgG and IgA by ELISA in B/HPIV3 empty-vector control immunized hamsters 21/22 days after challenge with the three SARS-CoV-2 strains (Fig. S2). The strongest responses were detected by IgA ELISA in three of five animals after WA1/2020 challenge, in the same order of magnitude as those on day 21 after immunization with B/HPIV3/S-6P (Fig. S2B, top row). Overall, after SARS-CoV-2 challenge of B/HPIV3 empty-vector control immunized animals, primary S-specific nasal antibody responses were detected in fewer animals than on day 21 after immunization with B/HPIV3/S-6P and B/HPIV3/S-Delta-6P. In NW from the B/HPIV3/S-6P and B/HPIV3/S-Delta-6P-immunized hamsters, no or very weak anamnestic antibody responses were detected after SARS-CoV-2 challenge (Fig. S2A and S2B, middle panels), with small post-challenge anti-S IgG and IgA increases only in a few animals that had low or background levels of anti-S IgG or IgA three weeks after immunization. B/HPIV3/S-Omicron-6P-immunized hamsters had no to low anti-Wuhan-Hu-1 S IgG and IgA detectable in the upper airways three weeks following immunization; most animals in the B/HPIV3/S-Omicron-6P immunized groups exhibited anamnestic increases in nasal anti-S IgG and IgA after WA1, Delta, or BA.1 challenge, detectable by Wuhan Hu-1 S ELISA (Fig. S2A and S2B, right panels). The absence of post-challenge increases in most animals with nasal IgG and IgA detectable after immunization with B/HPIV3/S-6P or B/HPIV3/S-Delta-6P suggested that protection against SARS-CoV-2 challenge (Fig. 4) was restrictive with respect to anamnestic nasal antibody responses.

We also evaluated the mucosal antibody levels in the lower airways using BAL samples collected at necropsy (day 23/24 pc, Fig. 5A). Even though there was variability within the groups, anti-S IgG and IgA antibodies were detected in the lower airways of most of the B/HPIV3 empty vector-immunized hamsters 3 weeks after SARS-CoV-2 challenge, revealing primary mucosal antibody responses following high levels of challenge virus replication in these control animals (Fig. 4). Since it is not possible to obtain sequential BAL samples from the same animals, the kinetics of mucosal responses in the lower airways could not be assessed directly. Compared to the B/HPIV3 empty-vector control groups, post-challenge anti-S IgG and IgA titers in B/HPIV3/S-6P immunized animals were significantly higher in all challenge groups, reflecting either a strong and durable primary response in the lower airways to the B/HPIV3-vectored S-6P antigen, and/or a boost of mucosal antibodies after challenge. Compared to the post-challenge titers in B/HPIV3-empty vector controls, BAL anti-S IgG titers were higher in six of nine challenge groups that had previously been immunized with S-expressing B/HPIV3, and anti-S IgA titers were higher in four of nine of these challenge groups. Even though we were not able to directly compare pre- and post-challenge antibody levels in the BAL from the same animals, these results indirectly suggest that anamnestic mucosal antibody responses occurred in the lower airways in six of the nine groups immunized with the S-expressing B/HPIV3 vectors. In addition, we evaluated the serum IgG and IgA responses to the ancestral Wuhan-Hu-1 S and to the RBD on day 23/24 pc by ELISA (Fig. 5B and 5C, left panels; anti-RBD titers are shown in Fig. S3). We detected high anti-S and anti-RBD serum IgG and IgA titers in all challenged hamsters. After BA.1/Omicron challenge, B/HPIV3 empty-vector immunized control animals had about 10-fold lower Wuhan-Hu-1 S-specific IgG and IgA antibody titers than animals primed with any of the S-expressing versions of B/HPIV3, including the B/HPIV3/S-Omicron-6P primed animals (Fig. 5B and 5C, left panels). Comparison of paired post-immunization and post-challenge sera (Fig. 5B and C, right panels) revealed that the anamnestic responses to the S antigen after challenge in all groups that had previously been immunized with S-expressing versions of B/HPIV3 were limited (medians <16-fold), suggesting strong protection of the S expressing B/HPIV3 vectors against matched or heterologous challenge with these three variants. We noted that matched challenge of B/HPIV3/S-Omicron-6P primed animals induced the strongest anamnestic responses, detectable by Wuhan-Hu-1 S ELISA, possibly reflecting affinity maturation and increase of antigenic breadth after challenge. The comparison of the RBD-specific responses of the paired post-immunization and post-challenge sera showed that the anamnestic IgG and IgA responses to the RBD region after challenge in animals previously immunized with S-expressing B/HPIV3 vaccines were generally stronger than the anamnestic responses to the whole S protein, especially in the B/HPIV3/S-Delta-6P and B/HPIV3/S-Omicron-6P immunized groups, irrespective of the challenge virus (medians up to 79-fold) (Fig. S3).

To evaluate further the breadth of antibody responses post-challenge, sera were also evaluated for their ability to inhibit the binding of soluble ACE2 protein to S of 20 SARS-CoV-2 variants (Fig. 5D and 5E). As shown in the heat maps (Fig. 5D) and radar plots (Fig. 5E), sera from B/HPIV3 empty-vector immunized/WA1/2020 or B.1.617.2/Delta challenged hamsters efficiently inhibited the binding of ACE2 to S of Wuhan-Hu-1 or B.1.617.2/Delta as well as early variants such as B.1.1.7 or B.1.351 (95 to 100% inhibition) (Fig. 5D, top and middle panel, top rows and Fig. 5E, left and middle panel, red lines). However, using sera from these animals, ACE2 binding inhibition to S from more recent variants was overall lower (44 to 85% inhibition). Sera from B/HPIV3 empty vector immunized/BA.1/Omicron challenged hamsters exhibited a different breadth with strong ability to inhibit ACE2 binding to S of BA.1, BA.2 and BA.3 variants (83 to 96% inhibition) but reduced ability to inhibit ACE2 binding to S from early or more recent SARS-CoV-2 variants including XBB variants (31 to 54% inhibition) (Fig. 5D, bottom panel, top row and Fig. 5E, right panel).

Sera from B/HPIV3/S-6P or B/HPIV3/S-Delta-6P immunized hamsters exhibited increased breadth after challenge with WA1/2020, Delta or BA.1/Omicron (Fig. 2D, 2E; 5D and Fig. 5E, blue and green lines) with almost complete ACE2 binding inhibition to all evaluated S antigens (77 to 100% inhibition). Finally, sera from B/HPIV3/S-Omicron-6P immunized hamsters also exhibited increased breadth after challenge with WA1/2020, Delta or BA.1, with nevertheless a lower ability to inhibit ACE2 binding to S proteins from the more recent SARS-CoV-2 isolates (BF7, BN.1, BQ, XBB and derivatives; 62 to 85% inhibition, Fig. 2D, 2E; 5D and Fig. 5E, orange lines).

## Discussion

Next-generation SARS-CoV-2 vaccines that could prevent infection via the respiratory route and transmission are needed for all age groups, including the pediatric population [29, 30]. Live-attenuated B/HPIV3-vectored SARS-CoV-2 vaccines that can be administered intranasally to the pediatric population could provide mucosal and systemic immunity and protection against SARS-CoV-2 as well as against HPIV3, an important pediatric pathogen. B/HPIV3 expressing the S-6P version of the ancestral Wuhan-Hu-1 strain (B/HPIV3/S-6P) was previously evaluated in hamsters and non-human primates [25, 26] and is currently being evaluated in a phase I clinical study in adults (Clinicaltrials.gov NCT06026514). If shown to be safe, versions of B/HPIV3 with updated S antigen from circulating variants are needed for evaluation in children.

In the present study, we generated live-attenuated B/HPIV3-vectored intranasal vaccines expressing the S-6P version of Delta or B.1.1.529/Omicron and evaluated their replication, immunogenicity and protective efficacy against homologous or heterologous challenge viruses in hamsters. There is no animal model for HPIV3 or B/HPIV3 vectors that is similar in permissiveness for HPIV3 to that of humans. Hamsters are a preclinical model to evaluate the replication, genetic stability, and immunogenicity of B/HPIV3 vectors. They are semi-permissive for B/HPIV3; following intranasal inoculation, B/HPIV3 replicates to high titers in the nasal epithelium and in the lungs of hamsters, without any clinical signs or histological changes. Moreover, the level of replication of BPIV3 is similar to that of HPIV3 in hamsters [31]. The level of replication of B/HPIV3/S-Delta-6P and B/HPIV3/S-Omicron-6P in both the NT and lungs of hamsters was comparable to that of B/HPIV3/S-6P, suggesting that differences in the sequences of the S antigen do not have a measurable effect on B/HPIV3 replication. In our previous studies, B/HPIV3/S-2P did not cause weight loss in hamsters, and by day 7 pi, both B/HPIV3/S-2P and B/HPIV3/S-6P replicated only to a low or undetectable level in the respiratory tract [24, 25]. Based on these results, B/HPIV3/S-Delta-6P and B/HPIV3/S-Omicron-6P are also expected to be cleared from the respiratory tract by around day 7 pi.

Since induction of mucosal antibody responses at the site of infection is critical for preventing infection and shedding, we collected nasal washes to evaluate the levels of anti-S IgG and IgA in the upper airways. Despite the assay limitations that arise during the evaluation of dilute nasal wash samples, we found that B/HPIV3/S-6P and B/HPIV3/S-Delta-6P induced mucosal S-specific IgA antibody responses in the upper airways of most immunized hamsters. We only detected low levels of anti-S IgA in the upper airways of a subset of B/HPIV3/S-Omicron-6P-immunized hamsters, likely due to the lower sensitivity of the Wuhan-Hu-1 S-based ELISA for antibodies induced by the B.1.1.529 S antigen of B/HPIV3/S-Omicron-6P. Thus, those titers might not be completely representative of the anti-S antibody titers induced by B/HPIV3/S-Omicron-6P, or by BA.1/Omicron challenge.

All three S-expressing B/HPIV3 vectors also induced robust serum anti-S and anti-RBD antibody responses. While we were not able to evaluate the mucosal antibody responses to all three vaccine-matched antigens, we evaluated the serum antibody response in antigen-matched S ELISAs using ancestral, B.1.617.2/Delta and B.1.1.529/Omicron S protein or RBD regions; we found that B/HPIV3/S-6P and B/HPIV3/S-Delta-6P induced serum IgG and IgA titers in the same order of magnitude against the S or RBD derived from the ancestral Wuhan-Hu-1 or the Delta variant, reflecting their close antigenic relatedness [32]. In contrast, B/HPIV3/S-Omicron-6P induced lower serum IgG/IgA titers to S antigens from the ancestral or B.1.617.2/Delta variant, and this difference was increased for IgG/IgA titers to RBD, confirming the antigenic differences between these S antigens [33, 34].

The antigenic differences between the Wuhan-Hu-1, B.1.617.2/Delta and B.1.1.529/Omicron S were evaluated further and confirmed in an ACE2 binding inhibition assay. Indeed, serum from B/HPIV3/S-6P- or B/HPIV3/S-Delta-6P-immunized hamsters efficiently and similarly inhibited ACE2 binding to S of early variants of concern such as B.1.1.7/Alpha or B.1.351/Beta but exhibited reduced ability to inhibit ACE2 binding to S antigen from BA.1/Omicron variants and derivatives. Interestingly, sera from these immunized groups showed greater inhibition of ACE2 binding to the S proteins of BA.4/BA.5, BA.2.75 and recently circulating XBB variants than sera from B/HPIV3/S-Omicron-6P-immunized hamsters. These sublineages originate from BA.2/Omicron, hence the divergence in antibody responses to pre-Omicron S and BA.1 S seems to be attributable to mutations that are not found in BA.2. For example, S proteins of BA.4/BA.5 and their sublineages including BA.4.6, BQ.1, BQ.1.1, BF.7 contain the mutation F486V. BA.2.75-derived sublineages including BA.2.75.2, BN.1, XBB.1, XBB.1.5 contain F486S/P and/or F490S in S, while BA.2.75 does not include these mutations [35].

WA1/2020 or B.1.617.2/Delta, but not BA.1/Omicron, induces weight loss in hamsters [28]. Immunization of hamsters with the B/HPIV3-S-expressing vaccines prevented weight loss following challenge with WA1/2020 or B.1.617.2/Delta. As expected, no weight loss was observed in animals challenged with BA.1/Omicron. Furthermore, no or low expression of inflammatory/antiviral genes was detected in the lungs of the B/HPIV3/S-6P- and B/HPIV3/S-Delta-6P-immunized hamsters after challenge with WA1/2020 or B.1.617.2/Delta, indicating that these two S antigens were cross-protective, reflecting their antigenic relatedness. However, a moderate increased expression of these inflammatory/antiviral genes was detected in these two groups of immunized animals when challenged with BA.1/Omicron, suggesting incomplete protection. On the other hand, groups of hamsters immunized with B/HPIV3/S-Omicron-6P exhibited increased expression of the inflammatory/antiviral genes in the lungs following challenge with WA1/2020 or B.1.617.2/Delta, also suggesting partial protection. Evaluation of SARS-CoV-2 replication in the NT and lungs after challenge of the immunized hamsters confirmed this partial protection. These results are in line with previously-published studies showing that vaccination with mRNA-based or adenovirus-based vaccines encoding for S from the ancestral Wuhan-Hu-1 strain provided robust protection against SARS-CoV-2 Delta infection, but low protection against BA.1/Omicron infection [36]. In our study, hamsters immunized with B/HPIV3 expressing S from B.1.1.529/Omicron were fully protected against lung inflammatory response and virus replication following challenge with SARS-CoV-2 BA.1/Omicron. These results are consistent with previous studies showing that mRNA-based vaccines encoding for S from BA.1/Omicron had superior immunogenicity against BA.1/Omicron compared to mRNA encoding for S from the ancestral Wuhan-Hu-1 strain [37].

Following challenge with SARS-CoV-2, the hamsters immunized with the S-expressing vectors did not exhibit a strong anamnestic antibody response in the upper airways. It is possible that the mucosal anti-S antibodies induced by the intranasal immunization prevented SARS-CoV-2 replication in the upper airways and thereby prevented the induction of an anamnestic antibody response. Indeed, hamsters that exhibited increased levels of anti-S IgA in the upper airways after SARS-CoV-2 challenge had low titers of anti-S IgA in the nasal washes after immunization.

A robust anti-S IgG and IgA antibody response also occurred in the lower airways and blood of most hamsters immunized with the S-expressing B/HPIV3 vectors. Following challenge with any of the SARS-CoV-2 viruses, B/HPIV3/S-6P- and B/HPIV3/S-Delta-6P-immunized hamsters exhibited a remarkable increase in breadth of their serum antibody responses. Serum from these animals almost completely blocked ACE2 binding to S of the previously-circulating SARS-CoV-2 variants, as well as the more recently-circulating variants. Hamsters immunized with the B/HPIV3/S-Omicron-6P and challenged with any of the SARS-CoV-2 variants also exhibited robust increases of the breadth of their antibody responses. However, serum from these animals still did not fully block ACE2 binding to S of the more recently-circulating SARS-CoV-2 variants. This suggests that immunization with B/HPIV3 expressing S of the ancestral strain or Delta variant primed for a broader antibody response than immunization with B/HPIV3 expressing B.1.1.529/Omicron S.

In conclusion, this study provided evidence that intranasal immunization with the live-attenuated B/HPIV3 vector expressing updated versions of SARS-CoV-2 S induced an anti-S antibody response in the upper and lower airways as well as in the blood of immunized in hamsters. These immunized hamsters were efficiently protected from weight loss, lung inflammatory responses and SARS-CoV-2 challenge virus replication in the upper airways and lungs. The long-term immunogenicity and protective efficacy of these B/HPIV3-vectored vaccines are currently being evaluated. Guided by these study results, B/HPIV3 vectors expressing S antigens from recent variants will be generated for clinical studies.

## Materials and Methods

### Ethics statement

Hamster studies were approved by the Animal Care and Use Committee of the National Institutes of Allergy and Infectious Diseases. The animal experiments were performed following the Guide for the Care and Use of Laboratory Animals by the NIH.

### SARS-CoV-2 viruses

The SARS-CoV-2 USA-WA1/2020 virus was obtained through BEI Resources (cat # NR-52281) and was passaged twice on Vero TMPRSS2 cells. SARS-CoV-2 isolate hVoV-19/USA/MD-HP05285/2021 (Lineage B.1.617.2; Delta Variant) was contributed by Andrew S. Pekosz and was obtained through BEI Resources (cat# 55673). The SARS-CoV-2 isolate hCoV-19/USA/HI-CDC-4359259-001/2021 (lineage B.1.1.529; Omicron variant) was deposited by the Centers for Disease Control and Prevention and obtained through BEI Resources (cat# NR-56486) and was passaged once on Vero TMPRSS2 cells. All experiments with SARS-CoV-2 were conducted in Biosafety Level (BSL)-3 containment laboratories approved for use by the US Department of Agriculture and CDC.

### Generation of recombinant B/HPIV3 expressing SARS-CoV-2 spike protein

A cDNA clone encoding the B/HPIV3 antigenome expressing the 1,273 aa full-length version of the 6P-stabilized SARS-CoV-2 S protein derived from the ancestral Wuhan-Hu-1 strain (GenBank MN908947), prefusion-stabilized by six proline substitutions [23] to generate B/HPIV3/S-6P, was generated previously [25, 26]. For the present study, we generated versions of B/HPIV3 that express the S-6P stabilized full-length versions of S proteins derived from SARS-CoV-2 B.1.617.2/Delta (GISAID EPI_ISL_3066877, sequenced by Long J, Renzette N, Adams M, Omerza G, Kelly K, Li L, The Jackson Laboratory, Farmington, CT) and B.1.1.529/Omicron variants [GISAID EPI_ISL_6795833, sequenced by Strydom A. et al., ZARV/NHLS, Department of Medical Virology, University of Pretoria [38]]. The ORF encoding the S-Delta-6P and S-Omicron-6P open reading frames (ORFs) (aa 1-1,273) were codon-optimized for human expression and include six proline substitutions to stabilize S in the prefusion form [23]. In addition, the S1/S2 polybasic furin cleavage motif “RRAR” was ablated and substituted by the “GSAS” motif [22]. Each ORF was framed by nucleotide adapters containing the BPIV3 gene start and gene end signal sequences. The resulting sequences were synthetized *de novo* and inserted into the B/HPIV3 antigenome cDNA using a singular *Asc* I restriction site in the 3’ noncoding region of the N gene, placing the additional gene between the N and P genes in the B/HPIV3 antigenome plasmid. The sequences of the antigenomic cDNAs were confirmed completely by Sanger sequencing, and plasmids were used to transfect BHK21 cells, clone BSR T7/5 [39], to produce the recombinant B/HPIV3 vectors expressing the S-6P antigens derived from B.1.617.2/Delta and B.1.1.529/Omicron isolates. Virus stocks were grown in Vero cells, and viral genomes purified from recovered virus were sequenced in their entirety by Sanger sequencing from overlapping uncloned RT-PCR fragments, confirming the absence of any adventitious mutations.

### Replication, immunogenicity and protective efficacy of B/HPIV3 S expressing vectors in hamsters

Five-to six-week-old male golden Syrian hamsters (*Mesocricetus auratus*) (n = 184) were obtained from Envigo Laboratories (Frederick, MD). The experiments were conducted in BSL2 and BSL3 facilities approved by the CDC. The timeline of the procedures and sampling is described in Fig. 1B. Two days before immunization, serum was collected from each hamster. On day 0, 46 hamsters per group (four groups total) were immunized under isoflurane anesthesia with 5 log_10_ pfu/hamster of B/HPIV3, B/HPIV3/S-6P, B/HPIV3/S-Delta-6P or B/HPIV3/S-Omicron-6P. On days 3 and 5 pi, five hamsters per immunized group were necropsied and nasal turbinates and lungs were harvested. Tissues were homogenized, clarified by centrifugation, and aliquots were snap frozen in dry ice. Vaccine replication was evaluated by titration of clarified supernatants using an immunoplaque assay. On day 21 pi, 18 of the 36 remaining hamsters per immunized group were randomly picked and nasal washes were performed under isoflurane anesthesia using 200 μl of 1X PBS. Aliquots were snap frozen in dry ice for evaluation of the mucosal antibody response by ELISA. On days 24 and 25 pi, serum was collected from the remaining 36 animals per immunized group.

Between days 26 and 28 pi, hamsters were transferred to a BSL3 facility for challenge with SARS-CoV-2. On day 32 pi, the 36 hamsters per immunized group were subdivided into three subgroups of 12 hamsters each and intranasally inoculated under isoflurane anesthesia with 4.5 log_10_ TCID_50_/hamster of SARS-CoV-2 WA1/2020, B.1.617.2/Delta or BA.1/Omicron challenge virus. Hamsters were monitored for weight loss and clinical signs of SARS-CoV-2 infection for 12 days after challenge. On day 35 pi (day 3 pc), six of the 12 hamsters per subgroup were necropsied and nasal turbinates and lungs were harvested. Then, tissues were homogenized and aliquots were snap frozen in dry ice for subsequent titration of SARS-CoV-2 challenge virus by determination of the 50% tissue culture infectious dose (TCID_50_) in Vero E6 cells and evaluation of the inflammatory response by RT-qPCR. On days 53 and 54 pi (days 21 and 22 pc), nasal washes of the six remaining hamsters per subgroup were performed under isoflurane anesthesia using 200 μl of 1X PBS. Aliquots were snap frozen in dry ice for evaluation of the mucosal antibody response by ELISA. On days 55 and 56 pi (days 23 and 24 pc), the remaining animals were necropsied and serum was collected. Bronchoalveolar lavage was also done on each animal using 1 ml of 1X PBS and aliquots were snap frozen in dry ice for evaluation of the mucosal antibody response by ELISA.

### Dual-staining immunoplaque assay

Replication of B/HPIV3 and derivatives was evaluated from nasal turbinates and lung homogenates by a dual-staining immunoplaque assay as previously described [24, 25]. Briefly, sub-confluent monolayers of Vero cells in 24-well plates were inoculated with 10-fold serially-diluted samples and incubated for 2 h at 32°C. Then, cells were overlaid with 1 ml per well of Opti-MEM (Thermo Fisher) containing 0.8% methylcellulose (Sigma), 1% L-glutamine (Thermo Fisher), 2.5% penicillin-streptomycin (Thermo Fisher), 0.5% amphotericin B (Thermo Fisher) and 0.1% gentamicin (Thermo Fisher). After incubation for 7 days at 32°C, cell monolayers were fixed overnight with ice-cold 80% methanol followed by 1 h incubation with Odyssey Blocking Buffer. Then, the HPIV3 antigens and SARS-CoV-2 S protein were detected by dual-immunostaining of plaques using a rabbit anti-HPIV3 serum and a human anti-SARS-CoV-2 S monoclonal primary antibody (CR3022) and the IRDye 680RD donkey anti-rabbit IgG and IRDye 800CW goat anti-human IgG secondary antibodies (LiCor). Plates were scanned using an Odyssey Infrared Imager (LiCor) and staining for PIV3 proteins and SARS-CoV-2 S was visualized in green and red, respectively, generating yellow plaque staining when merged.

### IgG/IgA dual ELISA

Expression and purification of the SARS-CoV-2 S-6P and RBD antigens from the Wuhan-Hu-1 strain was previously described [24]. Plasmids encoding SARS-CoV-2 S-2P from B.1.617.2/Delta or B.1.1.529/Omicron were a kind gift from Dr. Peter Kwong and Dr. I-Ting Teng (VRC, NIAID, NIH). S proteins were expressed and purified as previously described [40]. IgG and IgA antibody titers against SARS-CoV-2 S or its receptor binding domain (RBD) in sera, nasal washes or BAL samples were evaluated by a previously-described ELISA assay designed to detect IgG and IgA by sequential reads in the same sample. Briefly, black 96-well plates (MaxiSorp, Thermo Fisher, cat #437111) were coated with 75 ng per well of purified S or RBD in 50 mM carbonate coating buffer, and incubated overnight at 4°C. Plates were washed three times and blocked with 250 µl DPBS containing 5% dry milk (W/V). Samples were serially diluted in DPBS with 5% dry milk and 0.2% IGEPAL CA-630 by eleven 3-fold dilutions, starting from 1:100 (sera) or 1:10 (nasal washes and BAL). After a 1-hour incubation, plates were washed, and 100 µl per well of secondary antibodies [goat anti-hamster IgG(H+L)-conjugated with HRP (Thermo Fisher, cat# PA1-29626, 1:10,000, and rabbit anti-hamster IgA conjugated with biotin (Brookwood Biomedical, sab3002a, 1:1,000)] in dilution buffer were added. Plates were washed and 100 µl per well of diluted Streptavidin-Europium (Perkin Elmer, cat# 1244-360), diluted 1:2,000 in DPBS+ 0.2% IGEPAL CA-630, was added. Plates were incubated for 1 h and washed. Fifty µl of Pierce ECL Substrate (Thermo Fisher, cat# 32106) per well were added, and plates were read using a Synergy Neo2 (BioTek) plate reader to collect IgG luminescence signals. Plates were washed, and 100 µl per well of Enhancement Solution (Perkin Elmer: 4001-0010) was added. Plates were read again using a program for time-resolved fluorescence (TRF; excitation 360/40; emission 620/40) to collect IgA data. Results were determined by (i) calculating the average reading from duplicate wells, (ii) subtraction of the average reading from blank samples, (iv) setting the cut-off value to the blank average plus three standard deviations, and IgG and IgA titers of each sample were determined in sequential reads by interpolating the sigmoid standard curve generated on Prism 9.0 as previously described [41, 42].

### SARS-CoV-2 neutralization assay

SARS-CoV-2 neutralizing antibody titers in hamster sera were determined by live SARS-CoV-2 virus neutralization assays, performed in a BSL3 facility. Heat-inactivated sera were 2-fold serially diluted in Opti-MEM and mixed with 100 TCID_50_ of SARS-CoV-2 WA1/2020, B.1.617.2/Delta or BA.1/Omicron. After incubation at 37 °C for 1 h, the mixtures were added to quadruplicate wells of Vero E6 cells in 96-well plates and incubated for four days. The 50% neutralizing dose (ND_50_) was defined as the highest dilution of serum that completely prevented cytopathic effects in 50% of the wells and was expressed as a log_10_ reciprocal value.

### ACE2 binding inhibition assay

The ability of hamster sera to inhibit binding of ACE2 to SARS-CoV-2 S was evaluated in an ACE2 binding inhibition assay (Meso Scale Diagnotics, MSD). V-PLEX SARS-CoV-2 Panel 25 (K15586U), Panel 27 (K15609U) and Panel 34 (K15693U) Kits (MSD) were used. Each kit contains 96-well, 10-spot plates coated with soluble ectodomains of S proteins from the wild-type SARS-CoV-2 (Wuhan-Hu-1) and variants (Alpha, Beta, Delta and Omicron sublineages) representing a total of 20 different S antigens from the three different kits. Each assay was performed according to the manufacturer’s instructions and as previously described [25]. Heat-inactivated hamster sera were diluted at 1:20 in MSD diluent and evaluated in duplicate. Plates were read using a MESO QuickPlex SQ 120MM. The average electrochemiluminescence signals from duplicate wells for each serum and in wells with diluent only were calculated. The ACE2 binding inhibition by each serum is expressed as percent inhibition relative to the diluent.

### Quantification of SARS-CoV-2 viral genomes and host gene expression in lungs

One hundred μl of lung homogenate from each of six hamsters per subgroup from the day 3 post-SARS-CoV-2 challenge time point was subjected to total RNA extraction using TRIzol LS Reagent and Phasemaker Tubes Complete System (Thermo Fisher) in combination with PureLink RNA Mini Kit (Thermo Fisher). The extracted RNA was used to evaluate the level of expression of inflammatory/antiviral host genes as well as to quantify SARS-CoV-2 viral genomes by RT-qPCR as previously described [25].

Briefly, total RNA was used to synthesize cDNA using High-Capacity RNA-to-cDNA Kit (Thermo Fisher). Then, the level of expression of 10 inflammatory/antiviral host genes (CCL2, CCL3, CCL5, IFN-G, IFN-L, IL-1B, IP-10, IRF7, MX2, TNF-A) was evaluated in triplicate by RT-qPCR using TaqMan Fast Advanced Master Mix (Thermo Fisher) and previously-described TaqMan assays [43–45] on the QuantStudio 7 Pro. Beta-actin was also included as a housekeeping gene. The relative level of expression of each evaluated gene was expressed as log_2_ fold changes over the expression determined from five non-immunized, non-challenged hamsters [25].

Quantification of SARS-CoV-2 viral genomes from total lung RNA was performed as previously described [25]. Briefly, SARS-CoV-2 subgenomic E and N mRNA was quantified in triplicate by RT-qPCR using TaqMan RNA-to-Ct 1-Step Kit (Thermo Fisher) and previously-reported TaqMan primers and probes [46–49] on QuantStudio 7 Pro Real-Time PCR System (Thermo Fisher). Standard curves were generated by serially diluting pcDNA3.1 plasmids containing each target gene sequence. The limit of detection was 5.1 log_10_ copies per g of tissue.

### Statistical analyses

Data sets were assessed for significance using one-way ANOVA with Tukey’s or Sidak’s post-test or two-way ANOVA with Sidak’s post-test using Prism 9.0 (GraphPad Software). Time course data sets were assessed by mixed-effects analysis with Dunnett post-test; exact p values are indicated for levels of significance p<0.05. Data were only considered significant at P≤ 0.05. Details on the statistical comparisons can be found in the figures, figure legends, and results.

## Acknowledgements

This research was supported by the Division of Intramural Research of the National Institute of Allergy and Infectious Diseases (NIAID), National Institutes of Health (NIH) and by the Vaccine Research Center, an intramural Division of NIAID, NIH. We gratefully acknowledge all contributors, i.e., the authors and their originating laboratories responsible for obtaining the specimens, and their submitting laboratories for generating the genetic sequence and metadata and sharing via the GISAID Initiative, from which the sequences of the S proteins of the B.1.617.2/Delta and B.1.1.529/Omicron variants were derived. The funders had no role in study design.

## Author contributions

Conceptualization: H.-S.P., Y.M., C.LN., U.J.B.. Formal analysis: H.-S.P., Y.M., C.S., C.L., X.L., L.Y., J.A.K., E.F.D, U.J.B., C.LN.. Funding acquisition: P.D.K., U.J.B.. Investigation: H.-S.P., Y.M., C.S., C.L., X.L., L.Y., J.A.K., E.F.D., C.LN.. Methodology: H.-S.P., Y.M., C.S., C.L., X.L., L.Y., J.A.K., E.F.D, U.J.B., C.LN.. Resources: R.F.J., I-T T., P.D.K., U.J.B.. Supervision: CL.N., U.J.B.. Visualization: H.-S.P., Y.M., C.S., C.L., X.L., L.Y., J.A.K., E.F.D, U.J.B., C.LN.. Writing – Original Draft: H.-S.P., U.J.B., C.L.N. Writing – Review and Editing: All authors.

## Supplemental figures

**Fig S1.**
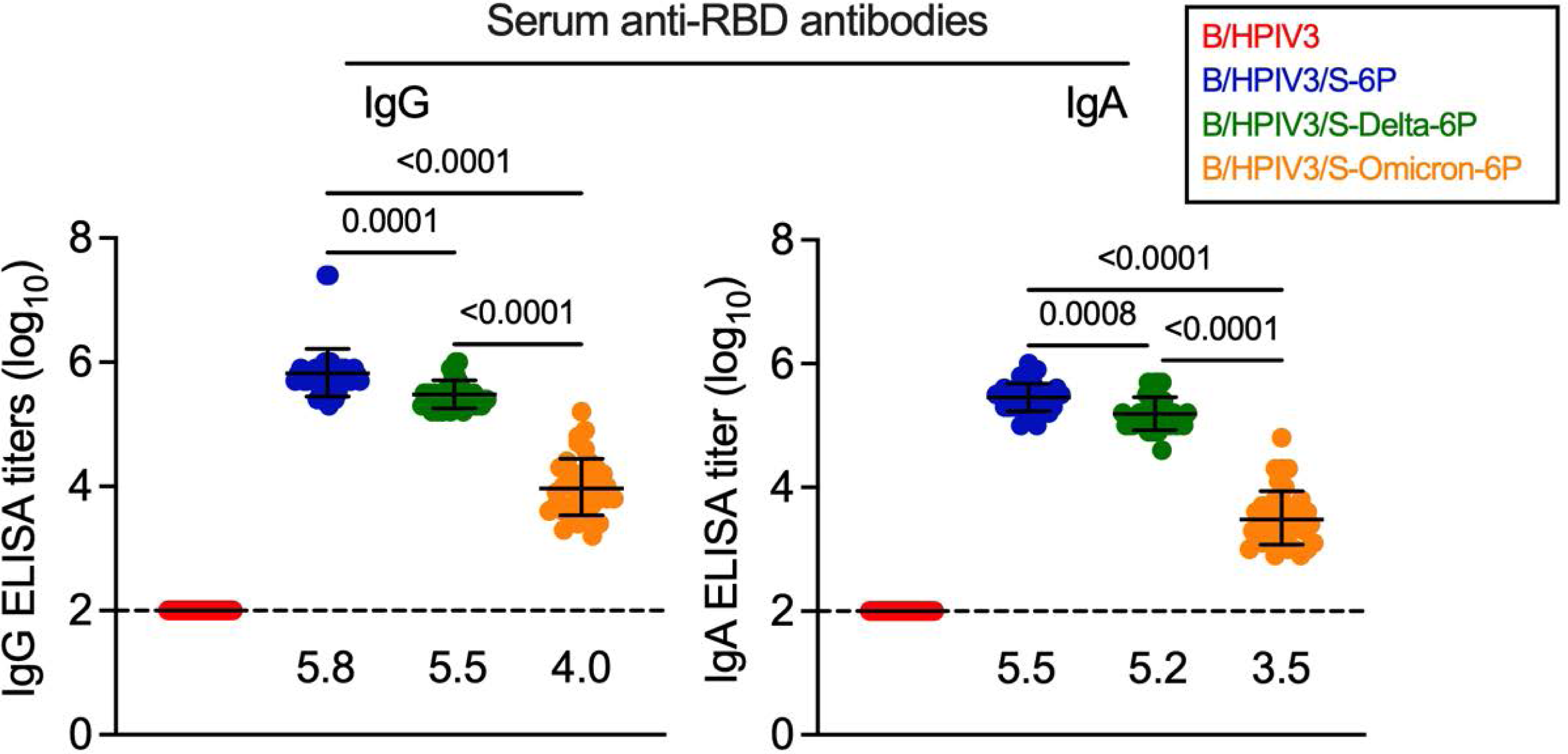
Serum antibody responses in immunized hamsters (related to Fig. 2B). On day 24 or 25 pi, serum was collected from n = 36 hamsters per group. Anti-RBD IgG (left panel) and IgA (right panel) serum antibody titers were evaluated by ELISA using purified RBD antigen preparations specific for the Wuhan-Hu-1 strain. IgG and IgA titers using purified S antigens matching the WA1/2020 or B.1.617.2/Delta or B.1.1.529/Omicron variants are shown in Fig. 2B. Each hamster is represented by a symbol, and medians with interquartile ranges are shown. The limit of detection (dotted line) is 2 log_10_. One-way ANOVA with Sidak post-test; exact p values are indicated for levels of significance p<0.05.

**Fig S2.**
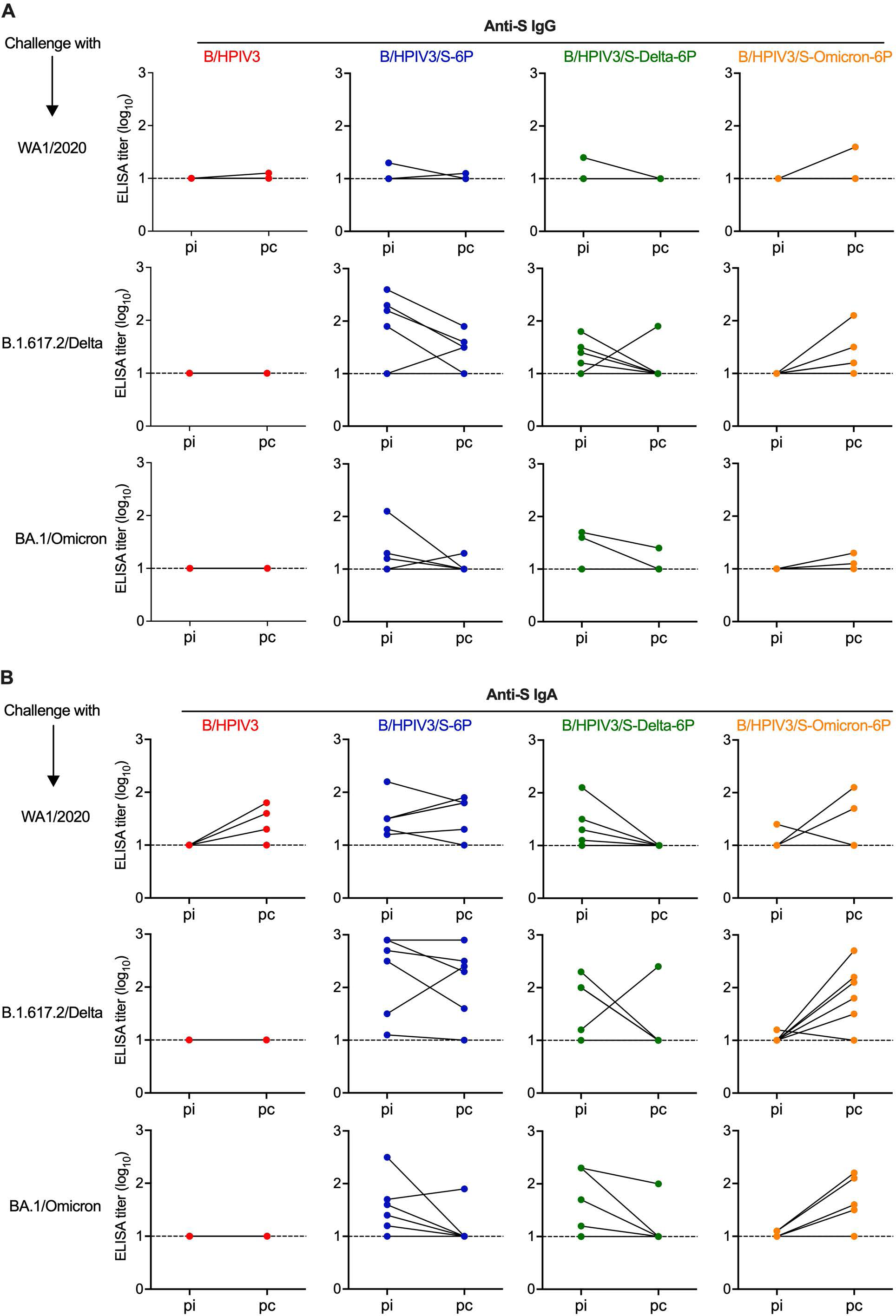
Anti-S IgG and IgA antibody response in upper airways of immunized and SARS-CoV-2 challenged hamsters (related to Fig. 2A). On day 21 post-immunization (pi), nasal washes were performed on 18 hamsters per immunized group, picked at random. On day 21/22 post-challenge (pc; equivalent to day 53/54 pi), nasal washes were performed on the six remaining hamsters per subgroup (see Fig. 1B for timeline of experiment). Anti-S IgG (A) and IgA (B) titers of paired nasal wash samples from the same six hamsters per subgroup were determined by ELISA. Note that ELISA titers from the samples collected on day 21 pi are also included in the results from 18 animals shown in Fig. 2A. Each hamster is represented by a symbol. The limit of detection of ELISA titers is 1 log_10_.

**Fig. S3.**
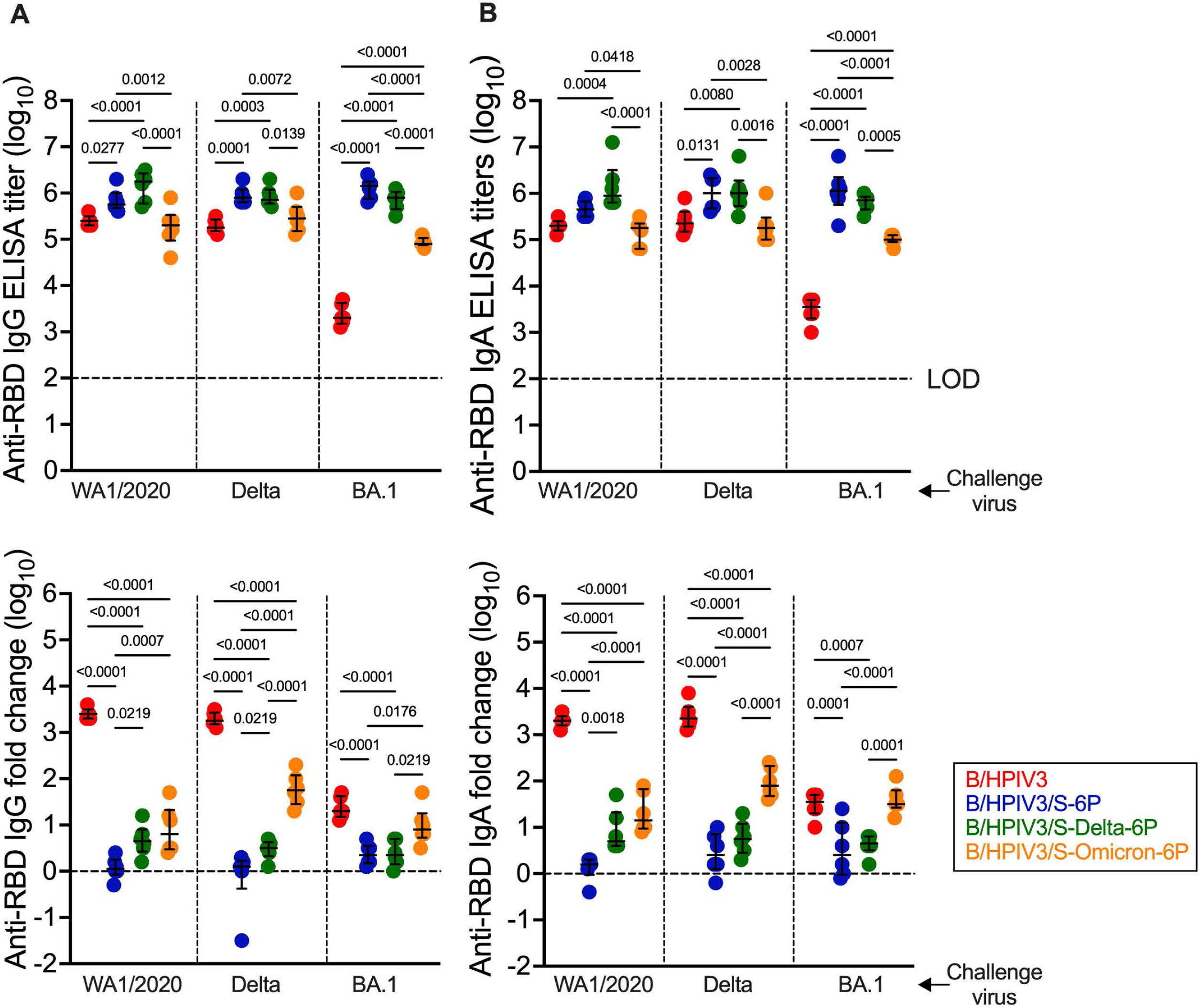
Serum anti-RBD IgG and IgA responses in immunized hamsters upon SARS-CoV-2 challenge (related to Fig. 5B and C). On day 23 or 24 pc, the six remaining hamsters per subgroup were euthanized, and sera were collected (see Fig. 1B for timeline of the experiment) for evaluation of the antibody response by ELISA. Serum anti-RBD IgG (A) and IgA (B) titers after challenge (top panels) and fold changes of post-challenge titers over post-immunization titers (bottom panels). N= 6 with the exception of n = 5 for B/HPIV3-immunized/WA1/2020 challenged. Purified preparations of RBD from the Wuhan-Hu-1 strain were used as an antigen in the ELISA assays. Each hamster is represented by a symbol and medians with interquartile ranges are shown. One-way ANOVA with Tukey post-test; exact p values are indicated for levels of significance p<0.05.

